# Cellular senescence is a central response to cytotoxic chemotherapy in high-grade serous ovarian cancer

**DOI:** 10.1101/425199

**Authors:** L. Calvo, S. Cheng, M. Skulimowski, I. Clément, L. Portelance, Y. Zhan, E. Carmona, J. Lafontaine, M. de Ladurantaye, K. Rahimi, D. Provencher, A.-M. Mes-Masson, F. Rodier

## Abstract

High-grade serous ovarian cancer (HGSOC) commonly responds to initial therapy, but this response is rarely durable. Understanding cell fate decisions taken by HGSOC cells in response to treatment could guide new therapeutic opportunities. Here we find that primary HGSOC cultures undergo therapy-induced senescence (TIS) in response to DNA damage induced by chemotherapy. HGSOC-TIS displays most senescence hallmarks including persistent DNA damage, senescence-associated inflammatory secretome, and selective sensitivity to senolytic Bcl-2 family inhibitors, suggesting avenues for preferential synergistic clearance of these cells. Comparison of pre- and post-chemotherapy HGSOC patient tissue samples revealed changes in senescence biomarkers suggestive of post-treatment “in patient” TIS, and a stronger TIS response in post-chemotherapy tissues correlated with better 5-year survival rates for patients. Together, these data suggest that the induction of cellular senescence in HGSOC cells accounts at least in part for beneficial cellular responses to treatment in patients providing a new therapeutic target.

**One Sentence Summary:** Cellular senescence is a central beneficial response to chemotherapy in high-grade serous ovarian cancer both *in vitro* and in patient.

## INTRODUCTION

Ovarian cancer remains the most lethal gynecological malignancy in the United States and Canada (Seiden, 2015). Among the different subtypes of the disease, high-grade serous ovarian cancer (HGSOC) is by far the most common, making up roughly two thirds of cases (Reid et al., 2017). Currently, initial treatment consists of a cytoreductive surgery followed by six cycles of a platinum agent- and taxane-based chemotherapeutic regimen (Network, 2017). Although primary response rates are high, frailty associated with multiple rounds of chemotherapy and eventual cancer cell drug resistance limit long-term five-year overall survival to around 40% (McGuire, 2009). Nevertheless, the high initial response rates suggest that a greater understanding of the molecular basis of the cellular responses involved could provide avenues to improve therapeutic approaches.

Cellular responses to stress like DNA damage induced by radio- or chemotherapy are termed cell fate decisions. These include repair or bypass of the damage, cell death (programmed or catastrophic), and cellular senescence (permanent growth arrest) (Rodier and Campisi, 2011). Senescent cells are characterized by several senescence-associated (SA) hallmarks that contribute to their biological functions. Among these, most prominent are the SA proliferation arrest (SAPA), SA apoptosis resistance (SAAR) and the micro-environmentally active SA secretory phenotype (SASP)(Coppe et al., 2008; Goldstein et al., 2005; Hayflick and Moorhead, 1961; Hernandez-Segura et al., 2018; Malaquin et al., 2016; Rodier and Campisi, 2011). The SAPA functions as a direct tumor suppressive mechanism and is largely mediated by two key tumor suppressor pathways: p53/p21^WAF1^ and p16INK4A/Rb (Beausejour et al., 2003; Campisi and d’Adda di Fagagna, 2007; Narita et al., 2003; Rodier et al., 2007), although many other pathways contribute a large degree of redundancy in this program (Gonzalez et al., 2016; Hernandez-Segura et al., 2018; Munoz-Espin and Serrano, 2014). It is thought that the stability of the SAPA is maintained to a great extent through SA chromatin remodeling, in particular through the formation of SA heterochromatin foci (SAHF) at the loci of proliferation-promoting genes (Narita et al., 2003; Zhang et al., 2007). Though SA chromatin remodeling remains incompletely understood, the downregulation of nuclear lamin B1 seems facilitate this process, and is also a senescence hallmark (Freund et al., 2012; Shah et al., 2013). Alternatively, the growth factors, cytokines, chemokines, extracellular proteases, and extracellular matrix proteins that compose the SASP mediate senescent cells functions in tissue repair (Coppe et al., 2008; Freund et al., 2010; Freund et al., 2011; Hoenicke and Zender, 2012; Krizhanovsky et al., 2008; Malaquin et al., 2016; Rodier et al., 2009). In the context of cancer, senescence is undoubtedly beneficial in its role as a barrier to malignant transformation in pre-neoplastic lesions (Campisi and d’Adda di Fagagna, 2007; Kuilman et al., 2010; Rodier et al., 2007), and therapy-induced senescence (TIS) in damaged cancer cells may also be advantageous by preventing tumor cell growth (Chang et al., 1999; Collado and Serrano, 2010; Gonzalez et al., 2016). However, alterations to tissue microenvironments caused by SASP must be evaluated contextually (Coppe et al., 2010). For example, in a mouse model of liver cancer, the microenvironment created by the presence of senescent cells was shown to promote the immune clearance of damaged cancer cells, suggesting that senescence could enhance the therapeutic response (Xue et al., 2007). In contrast, TIS was demonstrated to promote treatment resistance in a mouse lymphoma model by creating survival niches for residual cancer cells (Gilbert and Hemann, 2010; Rodier et al., 2009). Currently, whether TIS is an overall beneficial or detrimental process during human cancer treatment remains largely unknown.

A few studies have begun to shed light on the relevance of senescence in ovarian cancer. For example, evidence of a DNA-damage response (DDR) barrier, often associated with senescence, has been reported in hyperplastic fallopian tubes, suggesting that senescence acts as a tumor suppressive mechanism in this context to prevent the progression of pre-neoplastic lesions to HGSOC, perhaps applying a selective pressure responsible for the extremely high penetrance of p53 mutations in this disease (Jones and Drapkin, 2013). Also, the induction of senescence via manipulation of a variety of exogenous or endogenous factors after malignant transformation has been shown in immortalized cell lines derived from various ovarian cancer subtypes, demonstrating that ovarian cancer cells can be forced into senescence via drug exposure (Chang et al., 1999), growth factor overexpression (Bitler et al., 2011), steroid exposure (Diep et al., 2013), or select miRNA overexpression (Liu et al., 2014; Weiner-Gorzel et al., 2015). However, recent studies suggest that immortalized ovarian cancer cell lines likely do not reflect the full spectrum of human disease due to clonal derivation and low spontaneous immortalization rates (Ince et al., 2015; Verschraegen et al., 2003), highlighting the need for further research in this field.

Here, using human HGSOC primary cells, patient tissues, and their associated clinical data, we find that culture stress and chemotherapy-induced DNA damage trigger senescence in primary HGSOC cell cultures. Furthermore, HGSOC cells undergoing TIS are found to harbor persistent DDR signaling and express a SASP. These cells are then found to be selectively eliminated by ABT-263, a proven senolytic agent that inhibits Bcl-2 and Bcl-xL. Comparison of pre- and post-chemotherapy patient tissues reveals changes in biomarkers suggestive of senescence after treatment. Lastly, correlation of the expression of senescence markers in post-chemotherapy tissues with clinical data shows TIS to be associated with ameliorated clinical outcomes, suggesting TIS is a potential beneficial target for HGSOC improvement.

## RESULTS

### HGSOC primary cultures undergo culture stress-induced senescence

To define a human disease model adequate for the study of HGSOC therapy-induced cell fate decisions (TICFD), we first retrospectively reviewed data from the long-standing CRCHUM ovarian cancer tissue bank that established more than 30 ovarian cancer cell lines. Primary culture data reveals the vast majority of hundreds of primary HGSOC cultures did not spontaneously immortalize into cancer cell lines. Instead, following limited proliferation in culture for a few passages, growth slowed and eventually stopped, and cells adopted a flattened enlarged morphology without evidence of extensive cell death, suggestive of senescence. For example, out of 86 primary HGSOC cultures in the year 2000, only one culture generated an immortal cell line, while the remaining cultures ceased proliferation after about 4.6 passages (Fig. 1a-b). Following culture condition optimization (years 2006-2007), 15 HGSOC cell lines were established out of a total of 187 primary cultures, yielding a success rate of 8.02% consistent with previous observations (Ince et al., 2015; Verschraegen et al., 2003) reinforcing the idea that spontaneous immortalization of HGSOC primary cells is a relatively rare event.

**Figure 1:**
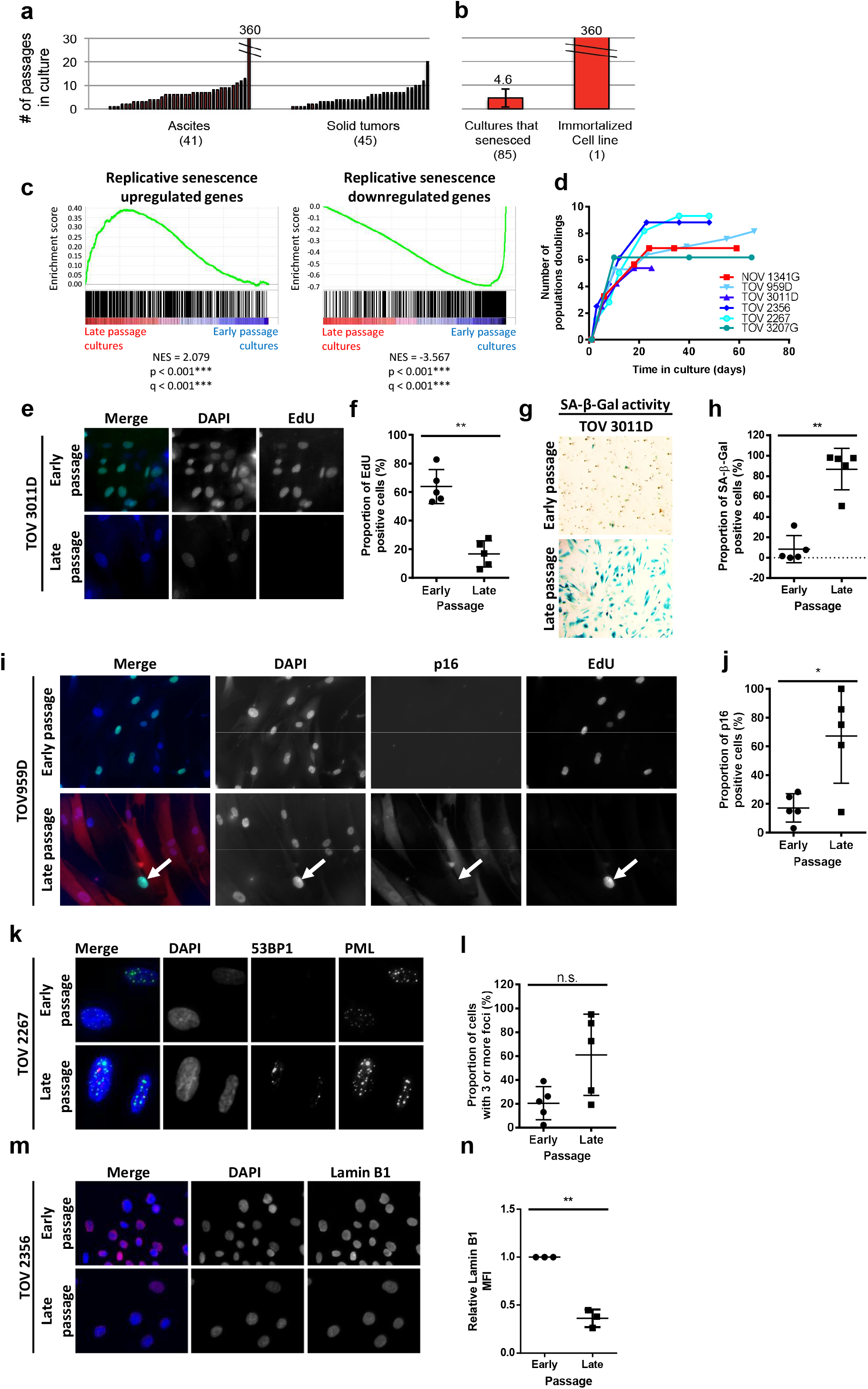
HGSOC primary cultures undergo culture stress-induced senescence. (**a**) The maximum number of passages before proliferation arrest of 86 independent HGSOC primary cultures collected during the year 2000. (**b**) The average maximal number of passages for growth-arrested cultures (85/86) or spontaneously immortalized cultures (1/86). (**c**) Gene set enrichment analysis (GSEA) plots of differentially expressed genes in late and early passage HGSOC cultures for genes found to be upregulated (left) and downregulated (right) in the normal human fibroblast replicative senescence gene expression signature (Hernandez-Segura et al., 2017). (**d**) Proliferation curves of normal epithelial ovarian primary cells (NOV1341G) and primary HGSOC cells (TOV959D, TOV3301D, TOV2356, TOV2267, and TOV3207G) serially passaged until proliferating arrest. Data is reported as population doublings (PD) over time. (**e**) Early- and late-passage HGSOC (TOV959D, TOV1294G, TOV2267, TOV2356, TOV3011D) primary cultures were labelled with EdU (24-hour pulse) (representative TOV3011D images shown). (**f**) Quantification of the percentage of EdU-positive nuclei from (e). (**g**) Early- and late-passage HGSOC (TOV959D, TOV1294G, TOV2267, TOV2356, TOV3011D) primary cultures were stained for SA-β-Gal activity (representative TOV3011D images shown). (**h**) Quantification of the percentage of SA-β-Gal positive cells from (g). (**i**) Early- and late-passage HGSOC primary cultures (TOV959D, TOV1294G, TOV2267, TOV2356, and TOV3011D) were stained for p16INK4A (red) and EdU (24-hour pulse; green) using immunofluorescence. Nuclei are counterstained in blue (DAPI) (white arrow highlights an EdU-positive, p16-negative cell) (representative TOV959D images shown). (**j**) Quantifications of p16INK4A-positive cells from (i). (**k**) Early- and late-passage HGSOC primary cultures (TOV959D, TOV1294G, TOV2267, TOV2356, and TOV3011D) were stained for 53BP1 and PML. DNA-SCARS are double-positive for both 53BP1 and PML-NBs (representative TOV2267 images shown). (**l**) Quantification of cells with more than three 53BP1 DNA damage foci from (k). (**m**) Early- and late-passage HGSOC primary cultures (TOV1294G, TOV2267, and TOV2356) were stained for lamin B1 (red) and nuclei counterstained with DAPI (blue) (representative TOV2356 images shown). (**n**) Quantification of the mean fluorescent intensity (MFI) of peripheral nuclear lamin B1 from (m) normalized to the control of each primary culture. Peripheral nuclear lamin B1 MFI was quantified within a 6 µm wide area along the perimeter of the nuclear area. Statistical significance was calculated using the paired t test for all panels except panel b, where the Kolmogorov-Smirnov statistic was used. n.s. not significant; * p < 0.05; ** p < 0.01; *** p < 0.001, **** p < 0.0001.

We further explored this apparent senescence response using Affymetrix gene expression analysis profiles from 42 independent HGSOC primary cultures at various passages within the culture lifespan (from early to late passages). Individual cultures were first assigned a proliferation status using unsupervised clustering for E2F-response gene activity (Fig. S1). Proliferating early passage cells (≤50% lifespan completed) were then compared to slow-proliferating late passage cells (≥75% lifespan completed), revealing significant gene set enrichment analysis (GSEA) to multiple senescence GSEA signatures (15/22 gene sets; Fig. S1d), including very significant enrichment for replicative senescence (Fig. 1c).

To further confirm key senescence hallmarks directly, we serially passaged five HGSOC primary cultures (TOV959D, TOV3011D, TOV2356, TOV2267, TOV3207G). Just like normal ovarian surface epithelial cells (NOV1341G) and as expected during culture stress-induced senescence HGSOC primary cultures gradually ceased proliferation (Fig 1d). When compared to early passage cells from their matched parental culture, late passage normal and cancer cells displayed increased senescence-associated beta-galactosidase activity (SA-β-Gal) and decreased DNA synthesis (Fig. 1e-h, Fig. S2a-b). Tumor suppressor genes driving senescence are often mutated in cancer (Gonzalez et al., 2016; Hanahan and Weinberg, 2011). Indeed, the well-known tumor suppressor *TP53*, a master mediator of damage-induced senescence, is mutated in >95% cases of HGSOC (Ahmed et al., 2010; Cancer Genome Atlas Research, 2011; Jones and Drapkin, 2013) (Fig. S2c). However, a quick analysis of The Cancer Genome Atlas (TCGA) large ovarian cystadenocarcinoma cohorts (currently 617 combined cases) reveal that p16INK4A (*gene CDKN2A*), another important mediator of SAPA and a senescence biomarker (Rodier and Campisi, 2011), is mutated in less than 5 % of HGSOC cases (Fig. S2c). We thus performed an immunostaining for p16INK4A concomitantly with a DNA synthesis assay on early and late passage HGSOC cells. Compared to early passage cells, a significantly higher proportion of late passage cells were positive for p16INK4A, and cells that stained positive for p16INK4A were DNA synthesis negative, corroborating the integrity of this SAPA pathway in HGSOC primary cells (Fig. 1i-j). We then explored the SA accumulation of permanent DNA damage foci termed DNA-SCARS (DNA segments with chromatin alterations reinforcing senescence)(Rodier and Campisi, 2011; Rodier et al., 2011). To this end, we co-stained for 53BP1 and PML using immunofluorescence (IF), the colocalization of which has been shown to mark DNA-SCARS (Rodier et al., 2011). Although DNA-SCARS were detected via colocaliztion, we quantified a non-significant increase in the proportion of cells that harbored three or more 53BP1 foci (Fig. 1k-l), perhaps owing to high levels of basal DNA damage associated with HGSOC (Jin et al., 2016; Wang et al., 2017). Lastly, we explored lamin B1 levels, shown to be downregulated in senescent cells (Freund et al., 2012), and found that late passage HGSOC cells strongly downregulate nuclear lamin B1 as compared to their early passage counterparts (Fig. 1m-n, Fig. S2d). Taken together, these data suggest that most if not all HGSOC primary cells retain the ability to undergo culture stress-induced senescence, and this senescence is associated with SAPA, SA-β-Gal, nuclear lamina alterations, the expression of p16INK4A, and perhaps the accumulation of further DNA-damage.

### HGSOC primary cultures undergo TIS in response to DNA damage and chemotherapy

We next tested whether HGSOC primary cells undergo senescence in response to DNA damage induced by therapy, a phenomenon termed therapy-induced senescence (TIS). Five different cultures of primary cells were treated with either 10 Gy ionizing radiation (IR), in order to induce senescence-associated DNA double-stranded breaks, or a combination of 10 µM carboplatin and 30 nM paclitaxel for a period of 12 hours (CP) to mimic one cycle of chemotherapy in patients (Heijns et al., 2008; Rodier et al., 2011; Sakai et al., 2001; Siddiqui et al., 1997; van der Vijgh, 1991; Wiernik et al., 1987). We then evaluated senescence hallmarks 8 days after the beginning of treatments. For both IR and CP, when compared to their controls, treated cells stained positive for SA-β-Gal (Fig. 2a-c), were largely positive p16INK4A, negative for DNA synthesis (Fig. 2d-f), exhibited increased DNA-SCARS (Fig. 2g-h, Fig. S3a), and expressed lower levels of nuclear lamin B1 (Fig. 2i-j, Fig. S3b), validating the presence of senescence hallmarks. Senescence induced by DNA damage has been well described to be associated with a SASP (Malaquin et al., 2015; Nakamura et al., 2008). Thus, we probed the secretome in three HGSOC primary cultures (TOV3011D, TOV2267, TOV2356) using multiplex immunoassay for a variety of SASP factors. Eight days after treatments, in all three primary cultures, a strong upregulation of SASP factors was detected, although specific factors differed slightly between treatments and individual cultures (Fig. 3a-c), consistent with previously observed context-dependent variations in SASP (Coppe et al., 2010; Coppe et al., 2008; Malaquin et al., 2016) Therefore, HGSOC primary cells seem to undergo TIS in response to DNA damaging therapies, including treatment with a standard chemotherapeutic regimen.

**Figure 2:**
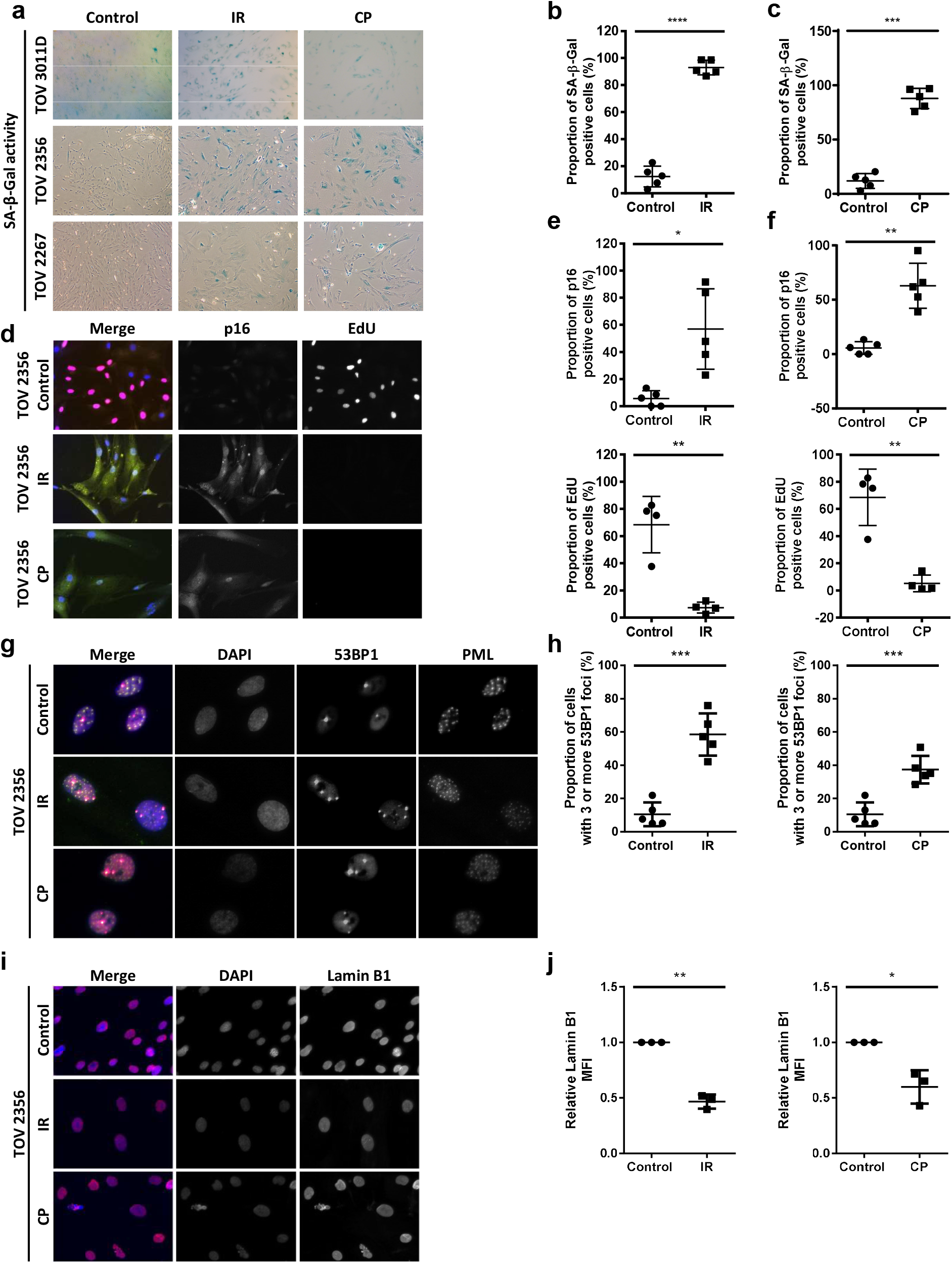
HGSOC primary cultures undergo TIS in response to DNA damage and chemotherapy. (**a**) HGSOC primary cultures were stained for SA-β-gal 8 days following treatment with 10 Gy ionizing radiation (IR) or a combination of 10 µM carboplatin and 30 nM paclitaxel for 12 hours (CP). (**b-c**) Quantification of the percentage of SA-β-gal-positive cells after IR (**b**) or CP (**c**) treatment from (a) in TOV 513, TOV1294G, TOV 2267, TOV 2356, and TOV3011D. (**d**) HGSOC primary cells were labelled with EdU (24-hour pulse; red) and stained for p16INK4A (green) 8 days after IR or CP treatment. (**e-f**) Quantification of p16INK4A-positive cells (top) and EdU-positive cells (bottom) in TOV513 (EdU staining not performed), TOV1294G, TOV2267, TOV2356, and TOV3011D cultures 8 days after treatment (e-f). (**g**) HGSOC primary cultures were stained for 53BP1 (red) and PML nuclear bodies (green) 8 days after IR or CP treatment. Colocalization of 53BP1 and PML (yellow in the merge panel on the left) reveals the presence of DNA-SCARS. (**h**) Quantification of cells with more than three 53BP1 foci in TOV513, TOV1294G, TOV2267, TOV2356, and TOV3011D cultures in either untreated conditions or after IR or CP treatment. (**i**) HGSOC primary cultures were stained for lamin B1 (red) 8 days after IR or CP treatment. (**j**) Quantification of peripheral nuclear lamin B1 MFI normalized to control in TOV1294G, TOV2267, and TOV2356 after IR or CP treatment. Peripheral nuclear lamin B1 MFI was quantified in a 6 µm wide area along the perimeter of the nuclear area. Statistical significance was calculated using the paired t test. * p < 0.05; ** p < 0.01; *** p < 0.001, **** p < 0.0001.

**Figure 3:**
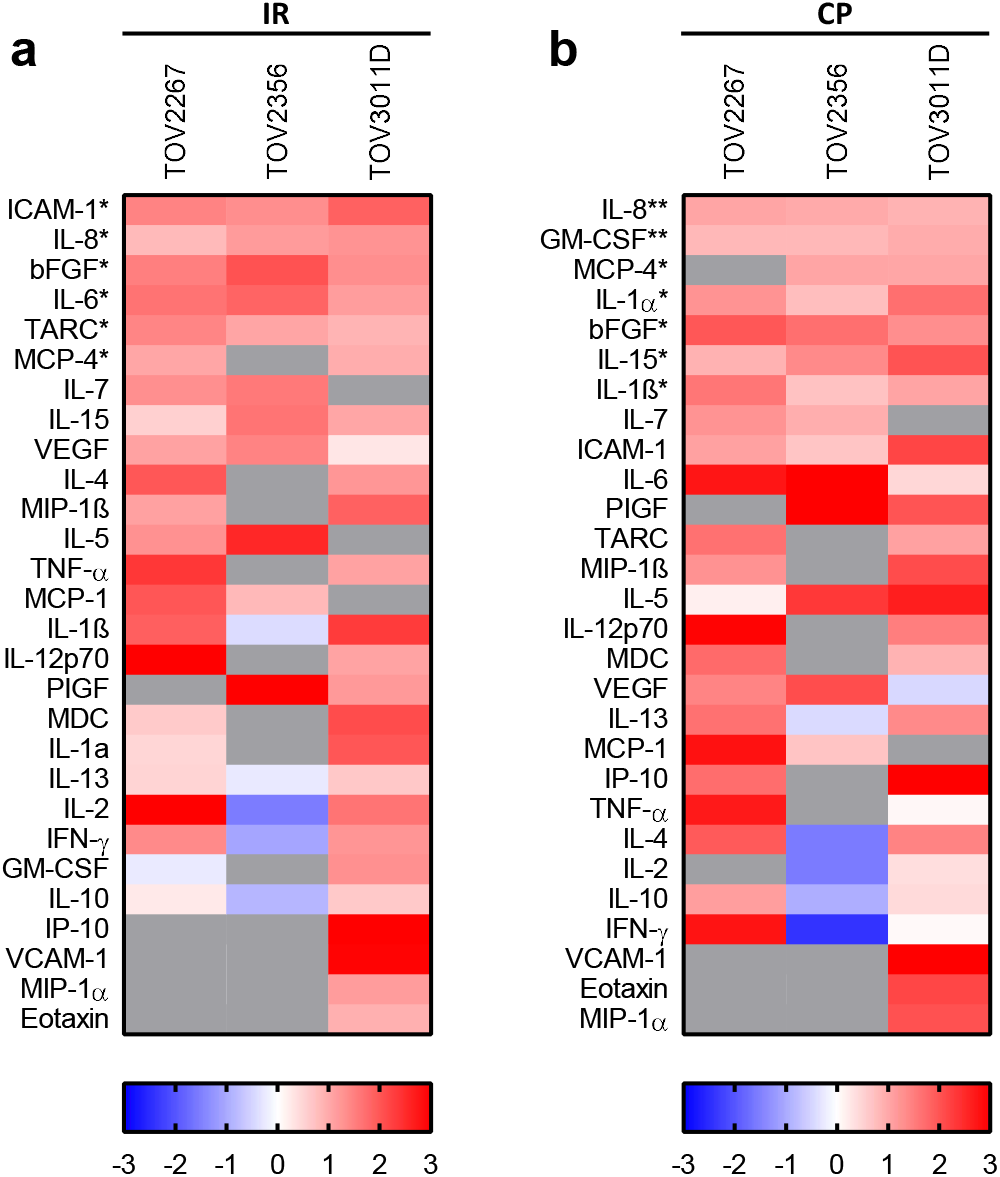
HGSOC primary cells trigger SASP secretomes in response to therapy. (**a-b**) Multiplex-based SASP profiling was performed on conditioned media from TOV2267, TOV2356, and TOV3011D HGSOC primary cultures treated (or not) with either 10 Gy IR (**a**) or carboplatin (10µM) and paclitaxel (30nM) (CP) for 12 hours (**b**). The conditioned medium was prepared 8 days following treatment (serum-free OSE medium for 16 hours). The heat-map colors indicate the relative fold change in protein levels (red for increased levels, blue for decreased levels relative to the control signal for each culture). Soluble factors with the highest fold changes analyzed are shown here. Factors are ordered, from top to bottom, according to increasing p-value. Gray: not detectable or not quantifiable; IR: ionizing radiation; CP: carboplatin and paclitaxel. Statistical significance was calculated using the paired t test. * p < 0.05; ** p < 0.01.

### HGSOC TIS renders cells sensitive to ABT-263

SAAR is a pharmacologically targetable senescence hallmark that represents an opportunity for treatment improvement (Chang et al., 2016; Wang et al., 2018). For example, senescent cells can be selectively re-directed to death using senolytic compounds like ABT-263 that inhibit the anti-apoptotic proteins Bcl-2 and Bcl-xL (Chang et al., 2016; Pan et al., 2017; Yosef et al., 2016; Zhu et al., 2016). Given the unpredictable microenvironmental impact of the SASP on tissue microenvironments and the possibility that HGSOC cell SAPA is not permanently stable, we sought to determine whether ABT-263 could be used to clear senescent HGSOC cells. We first treated pre-senescent TOV2267 cells with increasing concentrations of ABT-263 for 72 hours and assessed relative viability (Fig. 4a-b), revealing that pre-senescent HGSOC cells are not sensitive to ABT-263 concentrations up of 10 µM, much above the predicted active concentration of the compound (~0.1µM). Unlike their pre-senescent counterparts, similarly treated senescent TOV2267 cells (ABT-263 treatment from day 7-10 post-IR) showed decreased relative viability with concentrations of ABT-263 as low as 0.1 µM, with cells detaching and floating after ABT treatment (Fig. 4a, c). Similarly, pre-senescent TOV1294G and TOV513 cells were insensitive to ABT-263 at lower doses (Fig. 4d-e), but IR- or CP-treated senescent TOV1294G and TOV513 HGSOC cells were selectively preferentially eliminated by a 5 µM ABT-263 treatment (Fig. 4f-g). Thus, in addition to developing a TIS phenotype in response to therapy, HGSOC primary cells develop a sensitivity to ABT-263, suggesting a senolytic pharmacological strategy could enhance cancer cell elimination during current therapies.

**Figure 4:**
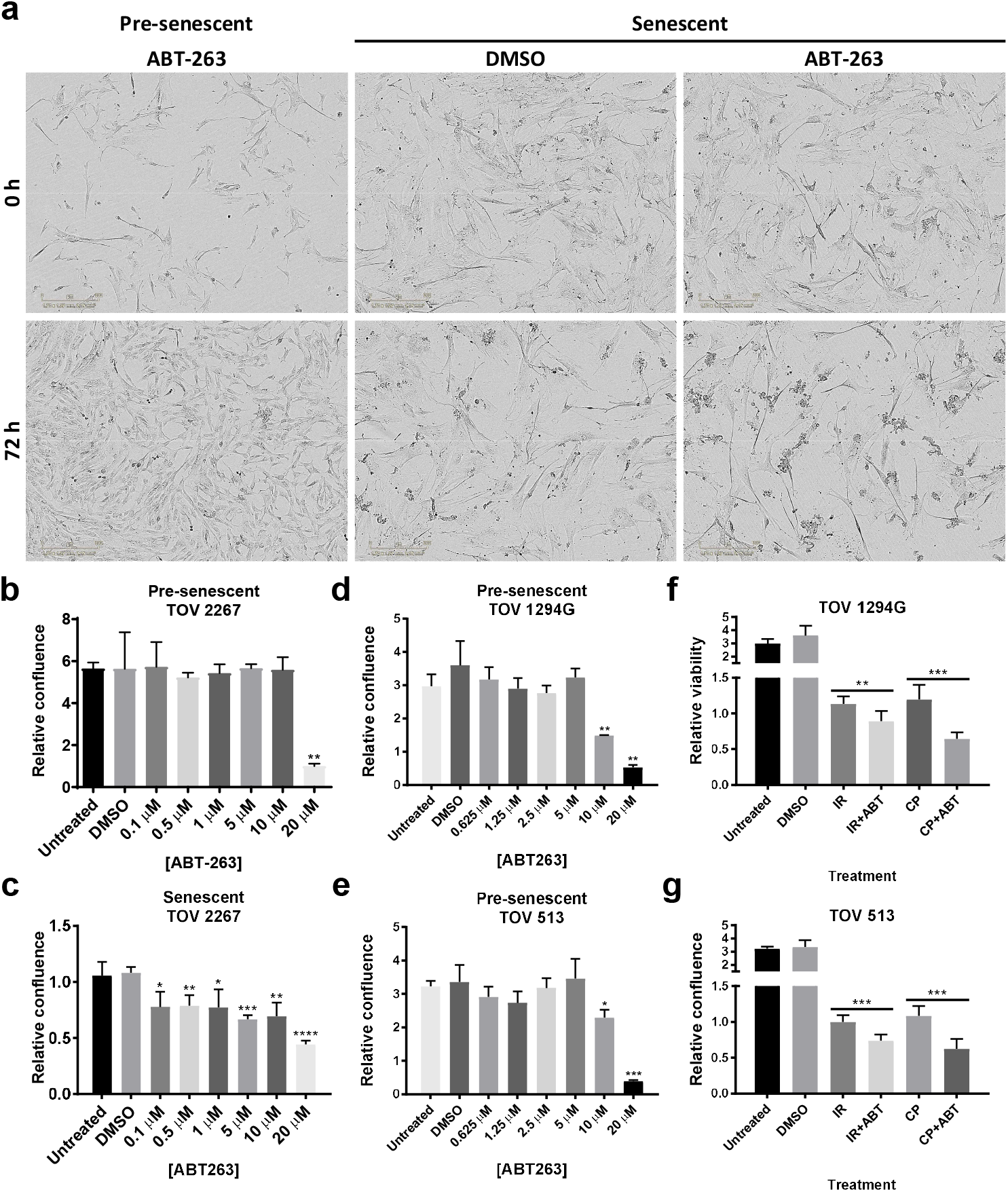
HGSOC TIS renders cells sensitive to senolytic clearance via ABT-263. (**a**) Presenescent or senescent (7 days after 10 Gy IR) TOV 2267 cells were treated with 5 µM of ABT-263 for 72 h. (**b-c**) Quantification of the relative confluence of pre-senescent (**b**) and senescent (**c**) TOV2267 cells when treated with different concentrations of ABT-263 for 72h. Data is represented as mean of the ratio of the confluence after 72h of treatment over the confluence at the beginning of treatment. (**d-e**) Quantification of the relative confluence of pre-senescent TOV1294G (**d**) and TOV513 (**e**) cells after treatment with different concentrations of ABT-263 for 72h. (**f-g**) Quantification of the relative confluence of TOV1294G (**f**) and TOV513 (**g**) cells subjected to 10 Gy IR or 10 µM carboplatin and 30 nM paclitaxel (CP) treatment, cultured for 7 days after treatment, and subsequently subjected to a 72-hour treatment with 5 µM of ABT-263. Statistical significance was calculated using the unpaired t test. * p < 0.05; ** p < 0.01, *** p < 0.001, **** p < 0.0001.

### HGSOC tissues display senescence hallmarks following exposure to chemotherapy in patients

Given that TIS occurs in HGSOC primary cultures in response to chemotherapy regimens used in the clinic, we explored whether TIS also occur in patients as a response to treatment. We constructed a tissue microarray (TMA) from HGSOC tissues collected from 170 patients treated at the Centre hospitalier de l’Université de Montréal (CHUM): 85 patients who had undergone surgery prior to chemotherapy, providing pre-chemotherapy samples, and 85 patients who had undergone surgery after primary chemotherapy, providing post-chemotherapy samples. Clinical follow-up data was associated with each patient sample (Fig. S4). The TMA was stained using IF to detect SA biomarkers, digitalized at high resolution, and quantitatively analyzed. Tissue cores were identified, segmented into epithelial (i.e. cancer cells) and stromal compartments using epithelial markers (cytokeratin CK7, 18, and 19), further segmented into nuclear and extra-nuclear compartments using DAPI (nuclear DNA signal), and categorized in specific compartments (i.e. total epithelial, epithelial nuclear, total stromal, and stromal nuclear; Fig. 5a). As expected for ovarian cancer (McCluggage et al., 2002) chemotherapy caused a significant reduction in the relative epithelial surface area within tumor tissues also reflected by the overall diminishment in total core mean fluorescent intensity (MFI) for cytokeratins or E-Cadherin (epithelial), and increased vimentin (stromal)(Fig. 5b). We first explored senescence biomarkers’ in tissue compartments using matched pre- and post-chemotherapy samples collected from the same patients, termed paired samples (limited availability). Interestingly, the MFI of epithelial nuclear laminB1 and Ki67 consistently decreased, while total epithelial IL6 consistently increased post-chemotherapy (Fig. S5a-f), consistent with changes expected in the context of HGSOC senescence based on primary culture results. Similar trends were seen in the stromal compartment as well, albeit less consistently (Fig. S5a-f). Trends in IL8, vimentin, and p16INK4A were less clear in both the epithelium and the stroma (Fig S5g-m). To gain a broader view, we examined SA biomarkers MFI variations in the cohort as a whole. Consistent with a TIS response to treatment in HGSOC epithelium, epithelial nuclear Ki67 and lamin B1 decreased significantly in post-chemotherapy samples, whereas total epithelial IL6 increased (Fig. 5f). There was no change in apoptosis levels as measured using caspase 3 activation (cleavage). Interestingly, total epithelial vimentin was found to increase post-chemotherapy (Fig. 5f), suggesting epithelial-to-mesenchymal transition (Huang et al., 2012; Lamouille et al., 2014), a phenomenon also linked to paracrine effects of the SASP (Malaquin et al., 2013; Parrinello et al., 2005; Sun et al., 2012). On the other hand, and unlike results observed in primary cultures, no significant differences were observed for epithelial nuclear p16INK4A or total epithelial IL8 between pre- and post-chemotherapy samples (Fig. 5f). Similarly, trends in the MFI of several SA biomarkers were consistent with senescence in the tumor stroma. In particular, stromal nuclear Ki67 and lamin B1 decreased, while total stromal IL8 increased (Fig. 5g). Unlike epithelial tumoral tissue, we observed slightly increased apoptosis rates in the stroma, but no significant difference was found in stromal p16INK4A, IL6, or vimentin between pre- and post-chemotherapy tissues (Fig. 5g). Overall, these findings offer support to the idea that TIS occurs as a response to treatment in HGSOC cells *in vitro*, and in patient, suggesting TIS is an important response to current HGSOC chemotherapy treatment.

**Figure 5:**
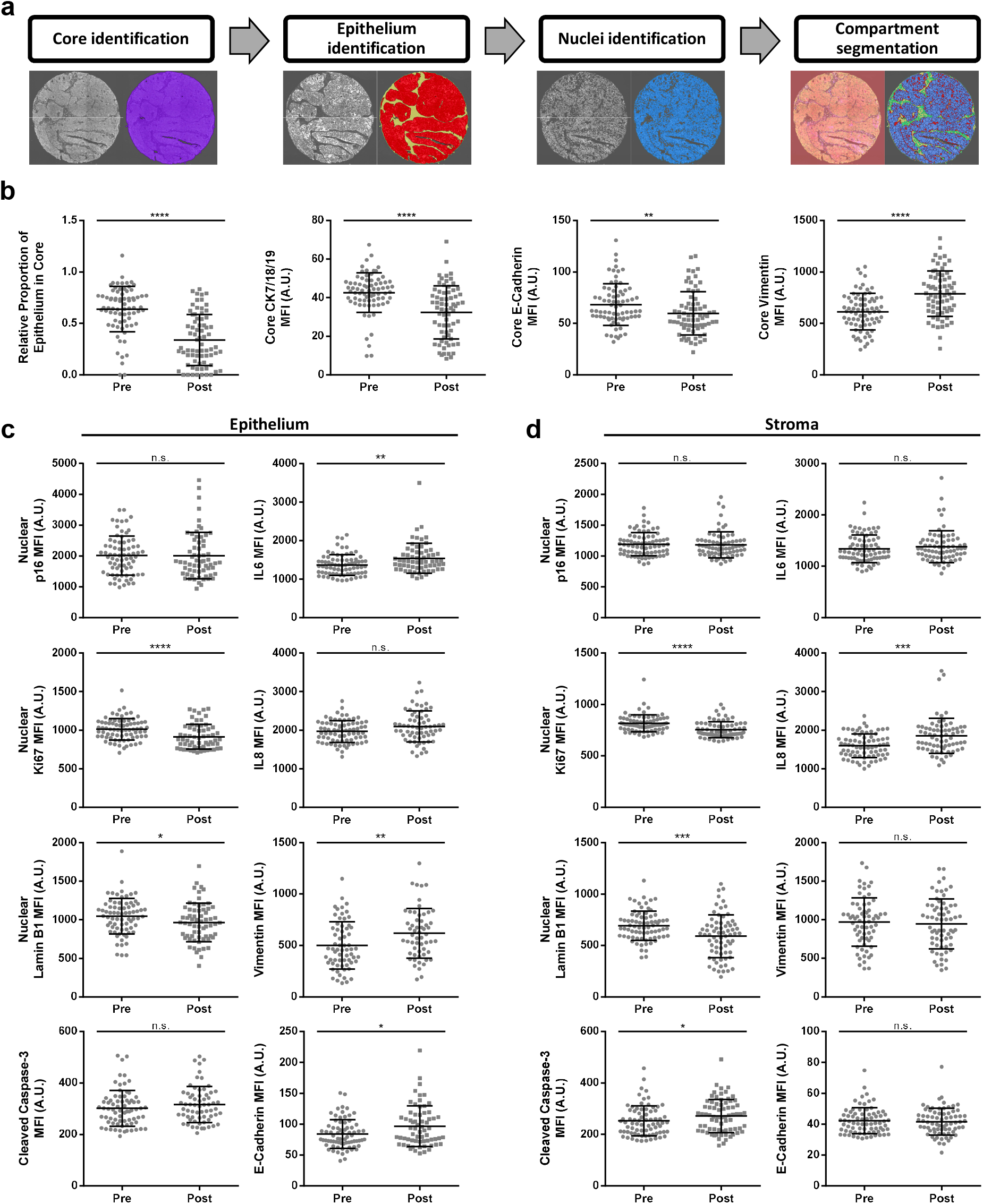
HGSOC tissues display senescence hallmarks following exposure to chemotherapy in patients. (**a**) Segmentation protocol of multi-channel IF images from a representative HGSOC tissue core. (**b**) Quantification of the proportion of epithelium in pre- and post-chemotherapy cores via the ratio of epithelium to core (far left), the total core MFI of the epithelial mask (cytokeratins 7, 18, 19) (middle left), the total core E-cadherin MFI (middle right), and the total core vimentin MFI (far right). (**c**) Quantification of the MFI of senescence markers (left, from top to bottom: nuclear p16, nuclear Ki67, nuclear lamin B, and cleaved caspase-3; right, from top to bottom: IL6, IL8, vimentin, and E-cadherin) in the epithelium of pre- and post-chemotherapy tissue cores. (**d**) Quantification of the MFI of senescence markers (left, from top to bottom: nuclear p16, nuclear Ki67, nuclear lamin B, and cleaved caspase-3; right, from top to bottom: IL6, IL8, vimentin, and E-cadherin) in the stroma of pre- and post-chemotherapy tissue cores. Statistical significance was calculated using the Mann Whitney test. n.s. not significant; * p < 0.05; ** p < 0.01, *** p < 0.001, **** p < 0.0001.

### Senescence-associated biomarker levels in post-chemotherapy HGSOC tissues correlate with 5-year survival

Whether senescence is beneficial or detrimental during human cancer treatment remains unknown. To determine how TIS may influence patient’s clinical course after treatment, we split patients from the post-chemotherapy group into high and low expressors of SA biomarkers and compared 5-year overall survival between the two groups using Kaplan-Meier analysis. Cutoff values for high- and low- expressors were determined via receiver operating characteristics (ROC) analyses. We find that low-expressors of epithelial nuclear lamin B1 post-chemotherapy, a sign of senescence, display improved 5-year overall survival as compared to high-expressors (Fig. 6a). Accordingly, high-expressors of epithelial nuclear p16INK4A, another sign of senescence, had improved survival in relation to low-expressors (Fig. 6b). Moreover, patients with low epithelial nuclear Ki67 MFI reflecting reduced proliferation trended towards improved 5-year overall survival, but the difference between the groups was not significant (Fig. 6c). While senescence-associated markers in the epithelium were predictive of 5-year overall survival, no marker in the stroma other than vimentin could significantly predict survival (Fig. 6d and Fig. S6a). Overall, a TIS response to chemotherapy appears associated with a positive prognosis for the patients.

**Figure 6:**
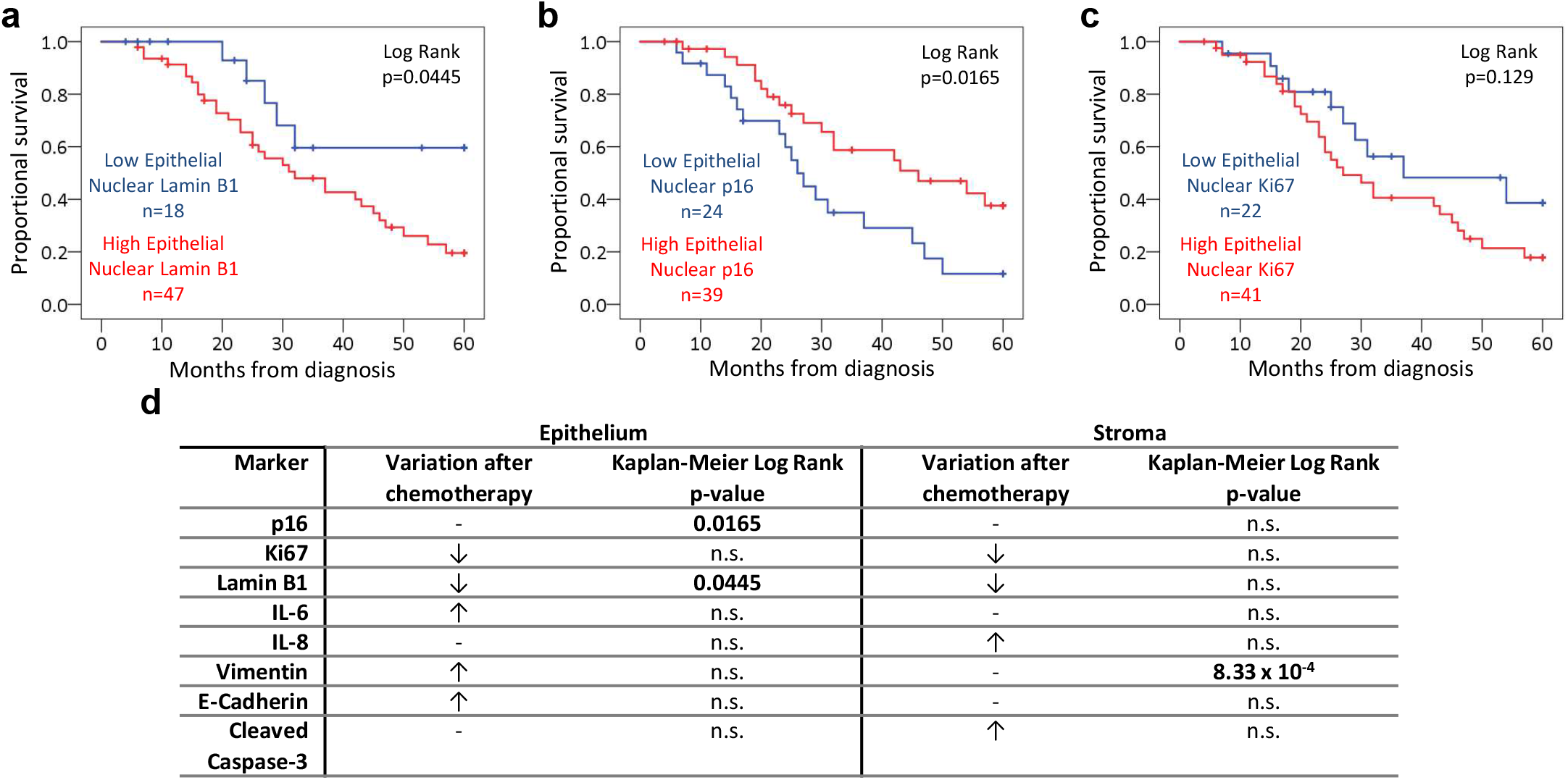
Senescence-associated biomarker levels in post-chemotherapy HGSOC tissues correlate with 5-year survival. (**a**) Kaplan-Meier analysis comparing 5-year survival between high and low expressors of lamin B1 after chemotherapy (n=65). Log rank p=0.0445. (**b**) Kaplan-Meier analysis comparing 5-year survival between high and low expressors of p16INK4A after chemotherapy (n=63). Log rank p=0.0165. (**c**) Kaplan-Meier analysis comparing 5-year survival between high and low expressors of Ki67 after chemotherapy (n=63). Log rank p=0.129. (**d**) Table summarizing the direction of variation of different senescence markers after chemotherapy in the epithelium and stroma, as well as their Kaplan-Meier log rank p-value comparing high and low post-chemotherapy expressors in terms of 5-year survival. Groups were separated into high and low expressors based on cut-offs determined by receiver operating characteristics (ROC) analyses.

## DISCUSSION

In this study we demonstrate that HGSOC primary cells taken from highly aggressive and fatal human cancers respond to chemotherapy by triggering TIS including most senescence hallmarks. Considering senescence is often driven by the p16/Rb and p53/p21 signaling cascades (Beausejour et al., 2003; Campisi and d’Adda di Fagagna, 2007; Narita et al., 2003; Rodier et al., 2007), and that TP53 is rendered dysfunctional through a multitude of mechanisms in nearly all HGSOC (Ahmed et al., 2010; Cancer Genome Atlas Research, 2011; Jones and Drapkin, 2013), it is perhaps not surprising that we found chemotherapy (CP) and IR promote p16INK4A-associated HGSOC senescence. Indeed, the key senescence tumor suppressor p16INK4A is rarely mutated in HGSOC (Fig. S2c), and its diffuse expression is used by pathologists to help differentiate HGSOC from other ovarian cancer subtypes (Kurman et al., 2014), raising the intriguing possibility that HGSOC cells maintain p16INK4A functions for cell cycle control in the absence of other TP53-associated DNA repair checkpoints, simultaneously preserving a capacity to undergo senescence in response to acute genome stress. A parallel non-exclusive possibility is that the p16/Rb pathway is inherently subverted downstream of p16INK4A in HGSOC, as often happen in HPV-induced cancers (Dyson et al., 1989; Sano et al., 1998; Ukpo et al., 2011), explaining low mutation rates of CDKN2A. Supporting both possibilities, our attempts to immortalize HGSOC cells through simple shRNA knockdown of p16INK4A failed in 2/3 primary culture tested (data not shown), suggesting that SAPA pathways other than the traditional p16/Rb and p53/p21 cascades remain to be identified in this context. In summary, the evidence presented here of p16INK4A-associated replicative senescence (>90% of primary cultures) combined with the low *CDKN2A* mutation penetrance in HGSOC patients (3-5%) suggests that the majority of HGSOC tumors retain the ability to undergo context-dependent TIS, providing treatment enhancement opportunities around pharmacological senescence manipulation.

In general, ethical and clinical considerations often preclude the collection of solid tumor tissue during chemotherapy regimens. HGSOC presents a particular clinical opportunity in that patients undergo either neoadjuvant chemotherapy followed by debulking surgery or primary debulking surgery followed by adjuvant chemotherapy (Kehoe et al., 2015; Vergote et al., 2010). These two treatment trajectories can provide access to either pre- or post- chemotherapy samples. By comparing tissues from patient cohorts that have received either treatment trajectories our data suggest that TIS occurs in ovarian cancer tissues in response to chemotherapy, and the associated clinical follow-up data further reveal that TIS may be beneficial in this context. This is important, as the activation of TIS and SASP following chemotherapy in HGSOC primary cultures and patient tissues suggest that ovarian cancer chemotherapy can alter post-treatment tumoral tissue microenvironments (Coppe et al., 2010; Gilbert and Hemann, 2010; Rodier et al., 2009; Xue et al., 2007), further guiding potential senescence pharmacological avenues toward SAPA reinforcement, SASP manipulation, and potential senescent cancer cell elimination via senolysis (Chang et al., 2016; Laberge et al., 2015; Moiseeva et al., 2013; Pan et al., 2017; Rader et al., 2013; Xu et al., 2015; Yosef et al., 2016; Zhu et al., 2016).

Of course, several observations were made that raise unanswered questions. For instance, why is p16INK4A upregulation not observed in TIS patient tissues, while it is readily observed in all TIS primary HGSOC cultures? It is important to note that post-chemotherapy tissues were collected a minimum of four weeks after the most recent chemotherapy cycle, as per standard clinical protocols to allow patients recovered. During this treatment black box, maybe cells that upregulate p16INK4A relative to the baseline expression indeed become senescent, but may have been subjected to immune clearance (Kang et al., 2011; Xue et al., 2007) or may have been lost by dilution to cell populations that have not efficiently activated TIS. In this context, it is also important to note that the post-treatment samples available for study are necessarily taken from patients with incomplete responses to chemotherapy (~75% of patients), thus it remains possible that tissues that have undergone a strong senescence response and subsequent immune clearance were underrepresented in this study. Conversely, it is possible that underrepresented tissues underwent cell fates other than senescence as well, such as apoptosis. Taken together, it is clear that a closer study of this immediate post-treatment period is paramount to improve our understanding of HGSOC TIS but it will require innovative clinical strategies to obtain he required human tissues.

Overall our dataset suggest that current chemotherapy regimens are efficient at inducing TIS in senescence-competent HGSOC cells, but that this TIS induction is probably suboptimal. It is likely that a subset of cancer cells either avoid or escape the senescence cell fate to potentially emerge from treatment. In light of this, the enhancement or reactivation of senescence in these cells may be an interesting and novel strategy in order to harness the curative potential of TIS in HGSOC. Perhaps even more important, the presence of biologically versatile senescent cells in a treated tumor implies the possibility that these influence other cells within the tumor in ways which alter treatment outcomes. For example, the DNA damage-induced secretion of IL6 in the senescent microenvironment of post-chemotherapy murine thymus is sufficient to create a protective niche for a subset of lymphoma cells that can cause cancer recurrence (Gilbert and Hemann, 2010; Rodier et al., 2009). Similarly, the presence of senescent cells in post-therapy murine breast cancers has been shown to trigger increased SASP levels which are associated with cancer recurrence (Jackson et al., 2012). Finally, the secretion of the SASP factor WNT16B (wingless-type MMTV integration site family member 16B) by senescent stromal cells in post-therapy murine prostates provides direct support for continued growth of prostatic epithelial cancer cells. Notably, increased stromal WNT16B levels have also been detected in post-therapy human prostate, ovarian, and breast cancers (Sun et al., 2012). Thus, senescence and its relationship with long-term treatment outcomes must be assessed in a context-dependent manner. Except for the emerging vision presented here for HGSOC, this concept has yet to be thoroughly studied in human cancers.

In the case of HGSOC, given the unpredictable influence of senescent cells on the clinical course, we further show that the incorporation of senolytic drugs into treatment regimens could prove to be beneficial via enhanced elimination of senescent cells. Several teams have reported the selective toxicity of compounds such as ABT-263 toward senescent cells (Chang et al., 2016; Pan et al., 2017; Zhu et al., 2016), and our results support the senolytic properties of this compound in the context of HGSOC. Research in this area is plentiful, and there exists a multitude of compounds with similar senolytic properties (Childs et al., 2017). Furthermore, elimimation of senescent cells by genetic manipulation proved to be effective in reducing aging-associated pathologies and tumor cells (Baker et al., 2011; Cheng and Rodier, 2015; Kohli et al., 2018; Laberge et al., 2013). Similarly, the use of a small molecule CDKi could be considered to strengthen p16INK4A-induced activities or epigenetic regulators to favor increased expression of p16^INK4A^ and other senescence genes (Bourdeau and Ferbeyre, 2016) prior to senolytic clearance. In summary, results this study warrant the idea that senescence is a therapeutic target for novel treatment modalities in HGSOC.

## MATERIALS AND METHODS

### Ethics statement

The CHUM institutional ethics committee approved this study. Informed consent from all patients was obtained before sample collection. Tumor and ascites samples were collected from patients following informed consent from the CHUM Department of Gynecologic Oncology.

### Patients and tissue specimens

Tumor samples were obtained from patients who underwent surgery for ovarian cancer in the CHUM Department of Gynecologic Oncology between 1993 and 2014. Disease stages were defined using the International Federation of Gynecology and Obstetrics (FIGO) staging system and tumor histopathology was classified using World Health Organization (WHO) criteria. Patient survival was calculated from the date of diagnosis, which corresponded to 3 months prior to surgery for post-chemotherapy samples. Five-year overall patient survival was defined as the time from diagnosis to death or last follow-up, up to a maximum follow-up of five years. Patients known to be alive at the time of analysis or 5 years after diagnosis were censored at the time of their last follow-up.

### Pre- and post-chemotherapy epithelial ovarian tumor tissue microarray (TMA)

A gynecologic pathologist reviewed all cases. The grade and type of ovarian carcinoma were identified and areas of interest were marked on hematoxylin and eosin–stained slides. Two cores of 0.8-mm diameter for each tissue sample were arrayed onto recipient paraffin blocks. The final TMA was composed of 340 cores from 170 epithelial ovarian tumors (2 replicate cores per patient). Clinicopathological details of the cohort are summarized in Fig. S4.

### Immunofluorescence staining of TMA

The TMA was sectioned into 4-µm thick slices and processed/stained using a BenchMark XT automated stainer (Ventana Medical System Inc., Tucson, AZ, USA). Antigen retrieval was carried out with Cell Conditioning 1 (Ventana Medical System Inc.; no. 950-124) for 60 minutes. The prediluted primary antibody was automatically dispensed and incubated for 60 minutes at 37°C. Primary antibodies used were mouse anti-p16INK4A monoclonal antibody (JC18) (Santa Cruz #sc56330), rabbit anti-Ki67 monoclonal antibody (SP6) (Thermo Fisher Scientific #RM-9106-S), rabbit anti-lamin B1 polyclonal antibody (Abcam #ab16048), rabbit anti-cleaved caspase-3 polyclonal antibody (Cell Signaling #6991), mouse anti-IL8 monoclonal antibody (6217) (R&D Systems #MAB208), rabbit anti-IL6 polyclonal antibody (Abcam #ab6672), mouse anti-vimentin monoclonal antibody (V9) (Sigma-Aldrich #V6630), and mouse anti-E-cadherin monoclonal antibody (G-10) (Santa Cruz #sc8426). The following steps were performed manually: After blocking with Protein Block serum-free reagent (Dako #X0909), Alexa-Fluor 568 secondary antibody (#A10037 Life technologies Inc., ON, Canada) and/or Alexa-Fluor 647 secondary antibody (#A31573 Life technologies Inc., ON, Canada (anti-rabbit); #A31571 Life technologies Inc., ON, Canada (anti-mouse)) were added for 45 minutes, followed by washing and blocking overnight at 4°C with diluted Mouse-On-Mouse reagent (Vector #MKB-2213). This was followed by an incubation of 60 minutes with an epithelial mask of 3 mouse monoclonal antibodies (anti-K7 clone OV/TL 12-30 (Thermofisher #MS-1352-P), anti-K18 clone DC-10 (Santa Cruz #sc6259), and anti-K19 clone ab-1 (Thermofisher #MS-198-P). The Alexa-fluor 750 secondary antibody (#A21037 Life technologies Inc., ON, Canada) was incubated for 45 minutes, followed by DAPI staining for 5 minutes. Staining with sudan black 0.1% for 15 minutes to quench tissue autofluorescence was followed by coverslip mounting using Fluoromount Aqueous Mounting Medium (#F4680, Sigma). The full TMA was scanned-digitalized using a 20X 0.75NA objective with a resolution of 0.325 µm (BX61VS, Olympus). Multicolor images were segmented and quantified using Visiopharm Integrator System (VIS) version 4.6.1.630 (Visiopharm, Denmark).

### VIS analysis

Immunofluorescence scoring was performed using the VIS software. Briefly, VIS was used to measure the MFI of all pixels in original digitalized multicolor TMA images for a chosen region of a tissue sample (total tissue core or core sub-compartments as described for Fig. 5a). Then, the MFIs corresponding to replicate total cores or selected core sub-compartments from the same patient are averaged to calculate one final mean MFI for each data point used in the presented analysis (See sFig. 7 for quality control correlation data on duplicate cores)

### Statistical analysis for TMA

Statistical significance of differences in MFI between pre- and post-chemotherapy groups was calculated using the Mann Whitney test. Survival curves were plotted using Kaplan-Meier analyses and the log-rank test was used to test for significance. All statistical analyses were done using the Statistical Package for Social Sciences software version 21 (SPSS, Inc.), and statistical significance was set at p<0.05. In a context-dependent manner patients were excluded from final analysis based on poor quality cores (folded or scratched cores) or for lack of epithelial material in cores. The breakdown of usable cores per analysis is as follow: Epithelial nuclear p16INK4A: pre- n=74, post- n=63. Epithelial nuclear Ki67: pre- n=76, post- n=63. Epithelial nuclear lamin B1: pre- n=73, post- n=65. Total epithelial cleaved caspase-3: pre- n=73, post- n=65. Total epithelial IL6: pre- n=75, post- n=64. Total epithelial IL8: pre- n=75, post- n=64. Total epithelial vimentin: pre- n=65, post- n=57. Total epithelial E-cadherin: pre- n=73, post- n=65. Stromal nuclear p16INK4A: pre- n=76, post- n=72. Stromal nuclear Ki67: pre- n=76, post- n=71. Stromal nuclear lamin B1: pre- n=75, post- n=71. Total stromal cleaved caspase-3: pre- n=75, post- n=72. Total stromal IL6: pre- n=75, post- n=71. Total stromal IL8: pre- n=75, post- n=71. Total stromal vimentin: pre- n=68, post- n=66. Total epithelial E-cadherin: pre- n=75., post- n=72.

### Primary cell culture

Associated clinical data were available for all HGSOC primary cells (TOV) and primary normal epithelial ovarian cells (NOV) used. The inclusion criteria used to select primary cells was their International Federation of Gynecology and Obstetrics (FIGO) classification and the subtype of ovarian cancer. All primary cancer cells used were HGSOC, with FIGO stage II and tumor grade ranging from 2 to 3. Primary cell cultures were derived from patient samples following procedures reported in our previous publications (Lounis et al., 1994). Unless otherwise indicated, cells were cultured in OSE media supplemented with 15% fetal bovine serum (FBS) and 1% streptomycin in a 5% O_2_ incubator. Proliferation was assessed using population doubling (PD) according to the following formula: last PD + (Log (final cell number) – Log (initial cell number plated) x 3.32).

### SA-β-gal staining

Senescence-associated β-galactosidase (SA-β-gal) was assessed as previously described (Laberge et al., 2013). Pictures were taken with a Nikon Eclipse TE300 microscope (Nikon Instruments Inc. Melville, NY, U.S.A).

### Irradiation and chemotherapy

Cells were seeded in appropriate vessels. 48 hours after seeding, cells were treated with either 10 Gy of X-irradiation (IR) using a Cesium 137 irradiator, or a combination of carboplatin (10 µM) and paclitaxel (30 nM). Carboplatin and paclitaxel were removed after 12 hours. Senescence hallmarks were tested 8-10 days later as indicated.

### Immunofluorescence staining of primary cell cultures

Cells were cultured in 4- or 8-well chamber slides (Nunc, Penfield, NY, USA). After treatment and incubation, cells were fixed in 10% formalin (Sigma) for 10 minutes and washed 3 times in PBS for 5 minutes at room temperature (RT) followed by incubation for 30 minutes in blocking buffer (1% IgG-free BSA, 4% donkey serum [Jackson ImmunoResearch, West Grove, PA, USA] in PBS), then by incubation with primary antibodies diluted in blocking buffer overnight at 4 °C. Cells were then washed and incubated with secondary antibodies and Hoechst for 1 hour at RT. Finally, slides were washed and mounted with Vectashield (Vector Labs, Burlingame, CA, USA) or ProLong® Gold Antifade reagent with DAPI (Life Technologies, Willow Creek Road Eugene, OR, USA). Images were acquired using a Carl Zeiss Axio Observer Z1 fluorescence microscope with AxioVision software (45 Valley brook Drive Toronto, ON, Canada) and presented with Photoshop CS (Adobe, San Jose, CA, USA). Images were then either analyzed visually or via computer-assisted quantitative image analysis. Computer-assisted quantitative image analysis was performed on raw images using the AxioVision ASSAYbuilder software (45 Valley brook Drive Toronto, ON, Canada), Physiology Analyst module. Cell nuclei were identified and delimited using the DAPI signal. For EdU and p16, the EdU and p16 signal mean fluorescent intensity in each cell nucleus were quantified. Cut-offs for positive and negative EdU and p16 staining were determined subjectively based on the lowest value that deviated from background noise in each cell culture. For lamin B1, the mean fluorescent intensity of the lamin B1 signal was quantified in a 6 µM-wide band along the periphery of the nucleus, stretching from the outer limit of the nucleus inwards. Unless otherwise specified in the figure legends, each data point presented represents the average value measured in one cell culture. Computer-assisted image analysis was used for cultures TOV1294G, TOV2267 and TOV2356 for p16, EdU and lamin B1 quantification. Quantification of the proportion of p16-positive and EdU-postitive cells in cultures TOV513, TOV959D and TOV3011D was performed visually. Quantification of the proportion of cells with 3 or more 53BP1 foci was performed visually for all primary cell cultures.

### DNA synthesis detection

Cells in chamber slides were pulsed with the modified thymidine analogue EdU for 48 hours before fixation. Cells were processed as per the manufacturer’s protocol to detect the incorporation of EdU into DNA using Click-iT chemistry (Invitrogen C10340) and Alexa fluor® 488 azide (No. A10266) or Alexa fluor® 647 azide (No. A10277). Following washing with PBS, immunofluorescence staining for p16INK4A was performed as described above. Finally, cells were washed with PBS, mounted with Vectashield (Vector Labs) or ProLong® Gold Antifade reagent with DAPI (Life Technologies, Willow Creek Road Eugene, OR, USA), and analyzed as for IF above.

### Real-time quantitative polymerase chain reaction (qRT-PCR)

RNA was extracted from cells or tissues using TRIzol reagent. Reverse transcription was performed using Invitrogen Superscrit III (#11752). qRT-PCR using Sybr green (Invitrogen, The SYBR® GreenER™ qPCR SuperMix kits) was performed with the following protocol: 2 minutes at 50°C, 10 minutes at 95°C, 40 cycles of 15 seconds each at 95°C, 30 seconds at 72°C, and 1 minute at 60°C. The primers used are listed in the table below.

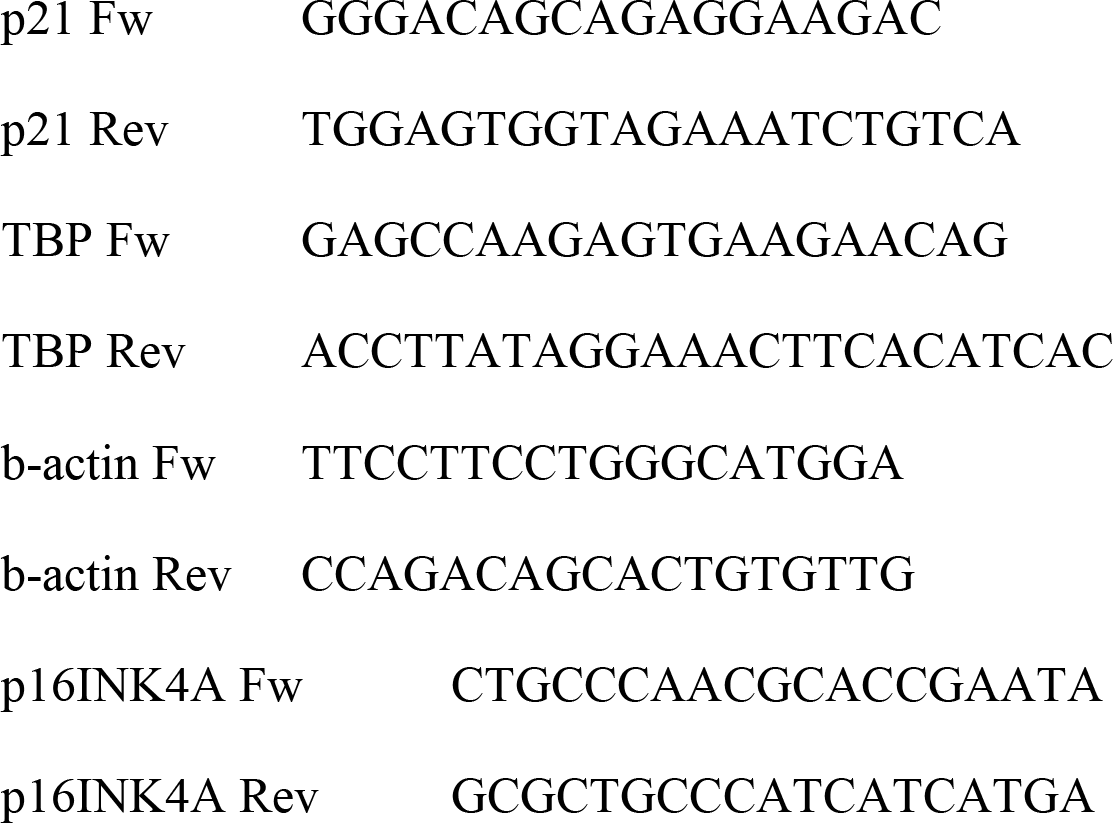

### Gene expression analysis

RNA from HGSOC cultures were extracted and purified as previously described (Gambaro et al., 2015; Zietarska et al., 2007) and microarray experiments were performed at the McGill University and Genome Quebec Innovation Centre (genomequebec.mcgill.ca) using HG-U133A GeneChip arrays (Affymetrix®, Santa Clara, CA). Gene expression levels were calculated for each probe set from the scanned image by Affymetrix® GeneChip (MAS5) as previously described (Cody et al., 2007). As reproducibility of expression data is variable at low raw expression values (Arcand et al., 2004), normalized data points below 15 were reassigned a threshold value of 15 based on the mean expression value of the lowest reliability scores. Probe sets having these threshold values in all samples were not used for further the analysis. Normalized gene expression data from 42 HGSOC primary cultures (sTable 1) was used to stratify expression clusters using unsupervised k-means (Gene Cluster 3.0) based on 177 E2F-target genes (Genes encoding cell cycle related targets of E2F transcription factors, M5925, http://software.broadinstitute.org/gsea/msigdb/cards/HALLMARK_E2F_TARGETS). Clustering results were generated using Java Treeview to highlight E2F-response clusters I, II, III; with cluster I reflecting high proliferation (high E2F-activity) and cluster II and III reflecting lower EF2-activiy (Fig. s1a-b). Early passage primary cultures from cluster I (≤50% replicative lifespan completed; n=5) were then compared to late passage primary cultures from cluster II-III (≥75% replicative lifespan completed; n=7) and the differential gene expression detected between these 2 groups (early versus late passage) compared using GSEA to 22 senescent signatures obtained from Hernandez-Segura’s work (Hernandez-Segura et al., 2017)(Fig. s1c-d).

### Antibodies and reagents

Antibody against 53BP1 (1:2000) is from Novus Biologicals Oakville, ON, Canada (No. NB100-304). Antibodies against PML (N-19) (1:500) goat (No. sc-9862) and p16INK4A (JC8) (1:100 or 1:300) mouse (No. sc-56330) are from Santa Cruz Biotechnologies (Santa Cruz, CA, United States). Antibody against γH2AX (ser 139) (1:2000) mouse (No. 05-636-I) is from Millipore (Temecula, CA, USA). Antibody against lamin B1 (1:1000) rabbit (No. Ab16048) is from Abcam (Cambridge, United Kingdom). The secondary antibodies used are all Alexa Fluor from Invitrogen (Carlsbad, CA, USA). Cy3 donkey anti-goat (1:800) from Jackson Immuno Research Laboratories (Baltimore Pike, West Grove, PA, USA) (No. 705-165-147), Alexa fluor 594 or 647 donkey anti-mouse for p16INK4A staining (1:800) (No. A1203 and No. A31571), Alexa fluor 488 (1:800) donkey anti-mouse for γH2AX (1:800) (No. A21202), and Alexa fluor 647 anti-rabbit (1:800) for 53BP1 and lamin B1 (No. A31573) are all from Life Technologies (Life Technologies, Grand Island, USA).

### SASP profiling

SASP was prepared as previously described (Rodier, 2013) with the following adjustments: Cells were seeded and treated with paclitaxel/carboplatin (or not) as described above, followed by incubation in OSE medium with 15% FBS for another 8 days (media was changed every 2 days). Sixteen hours before performing antibody array, the vessels were washed 3 times with PBS, and the media was replaced with serum-free media to generate serum-free conditioned media. Fresh conditioned media were collected and stored on ice for immediate use on multiplex ELISA, and cell numbers were counted. Multiplex secretome detection was performed according to the protocol provided by Meso Scale Discovery V-PLEX Human Biomarker (40-Plex Kit K15209D-2). In brief, 50-µl samples, diluted 1:2 or 1:4, were added to the wells. Plates were sealed and incubated for 2 hours at RT. After incubation, plates were washed 3 times with 150 µl of wash buffer per well, and 25 µl of detection antibody solution were then added to each well. The plates were incubated for another 2 hours at RT, followed by 3 times washing with wash buffer. Before reading, 150 µl of 2X read buffer T were added to each well. Finally, the plates were read on the MSD instrument, with signals normalized to cell number.

### Measurements of relative growth and viability

Cells were seeded in 96-wells plates and cultured within an IncuCyte® Zoom live cell imager that captured phase contrast images every 2 hours (Essen Bioscience). 24 hours after seeding cells received genotoxic treatments as indicated. Using captured images, the relative cell confluence (confluence mask) was determined by an integrated confluence algorithm (Essen Bioscience). To track treatment response, the data is reported as relative cell confluence (fold change) representing the confluence at the end of the indicated treatments over confluence just before treatment is initiated.

## ACKNOWLEDGEMENTS

We thank members of the Mes-Masson, Provencher and Rodier laboratory for valuable comments and discussions, as well as Suzana Anjos and Jackie Cheung for language editing. We thank the CRCHUM molecular pathology platform and the Institut du cancer de Montreéal (ICM) Imaging and Live imaging platforms. This work was supported by the ICM (DP, AMMM, FR) and by the Canadian Institute for Health Research (CIHR MOP114962 to FR), the Terry Fox Research Institute (TFRI 1030 to FR) and the Cancer Research Society (CRS) in partnership with Ovarian Cancer Canada (20087 to AMMM, DP; 22713 to FR). AMMM, DP and FR are researchers of CRCHUM/ICM, which receive support from the Fonds de recherche du Québec - Santé (FRQS). FR is supported by a FRQS junior I-II career awards (22624, 33070). Ovarian tumor banking was supported by the Banque de tissus et de données of the Réseau de recherche sur le cancer of the FRQS affiliated with the Canadian Tumor Repository Network (CTRNet). LCG and MS received ICM Canderel fellowships, SC received a FRQS postdoctoral award, MS received a CIHR doctoral award and YZ was supported by an ICM/MITACS postdoctoral fellowship.

## AUTHORS CONTRIBUTIONS

LCG, SC, and FR designed the study. LCG, SC, IC, MS, and JL performed experiments, YZ and EC performed bioinformatics analysis, KR revised all pathology samples for TMA construction, LCG, SC, MS, IC, and FR collected and analyzed data. LP and MD performed tissue banking and provided technical assistance. LCG, SC, MS, and FR wrote the manuscript. AMMM and DP provided technical support, biobank access, molecular pathology expertise, conceptual advice, and revised the manuscript.

---

**Supplemental figure 1: Primary HGSOC cultures proliferative lifespan transcriptome analysis.** (**a**) Major steps in the workflow of the analysis of transcriptome data from HGSOC primary cell cultures. Transcriptome data was collected via Affymetrix microarrays from 42 HGSOC primary cell cultures, each at a different passage with respect to total proliferative lifespan of individual cultures. Unsupervised clustering was then performed according to E2F target gene expression level, yielding 3 clusters (**b**). Using the eventual maximal passage number for each primary cell culture, cultures were selected from clusters 1 and 3, respectively, if the passage at which transcriptome data was acquired was 50 % or less and 75 % or more of the maximal eventual passage number of that culture (**c**). These cell cultures were considered to be “early passage” and “late passage” respectively. Late passage cultures were compared to early passage cultures on gene set enrichment analysis (GSEA) using gene sets generated by Hernandez-Segura et al. (2017) and downloaded from https://www.ncbi.nlm.nih.gov/geo/. (**b**) Unsupervised clustering of 42 HGSOC primary cell cultures according to E2F target gene expression. (**c**) Primary cell cultures included in the “early passage” (n=5) and “late passage” (n=7) groups. Primary cell cultures were selected from clusters I (high proliferation) and II-III (low proliferation) in (**b**), respectively, if the passage at which transcriptome data was acquired was 50 % or less and 75 % or more of the maximal eventual passage number of that culture, in order to form the early and late passage groups. (**d**) Gene sets described by Hernandez-Segura et al. (2017) significantly upregulated in “late passage” (left) and “early passage” (right) cell cultures on GSEA.

**Supplemental figure 2: Normal and HGSOC primary cultures undergo proliferative senescence.** (**a**) Early- and late-passage normal ovarian epithelial cells (NOV1341G) primary cultures were labelled with EdU (24-hour pulse). (**b**) Early- and late-passage normal ovarian epithelial cells (NOV1341G) primary cultures were stained for SA-β-Gal activity. (**c**) Mutation rates for TP53 and CDKN2A found in two large The Cancer Genome Atlas (TCGA) ovarian cystadenocarcinoma cohorts, including 311 cases (provisional) and 316 cases (Cancer Genome Atlas Research, 2011), using cBioportal. (**d**) Quantification of the relative mean fluorescent intensity of nuclear Lamin B1 per cell from (**Fig. 1 m**) (each cell represents a data point); statistical significance was calculated using the unpaired t test. **** p < 0.0001.

**Supplemental figure 3: HGSOC primary cultures undergo TIS in response to DNA damage and chemotherapy.**(**a**) Quantification of cells with more than three 53BP1 foci in TOV513, TOV1294G, TOV2267, TOV2356, and TOV3011D cultures in either untreated conditions or after IR or CP treatment (each high-powered field represents a data point). (**b**) Quantification of lamin B1 MFI per cell normalized to control in TOV1294G, TOV2267, and TOV2356 after IR or CP treatment (each cell represents a data point). Statistical significance was calculated using the unpaired t test. ** p < 0.01; *** p < 0.001, **** p < 0.0001.

**Supplemental figure 4: Clinical characteristics of the patients in the pre- and post-chemotherapy HGSOC cohorts.** (**a**) Clinical characteristics of pre-chemotherapy, post-chemotherapy, and both pre- and post-chemotherapy patients whose tumour tissue samples were used to construct the tissue microarray. (**b**) Cox regression analysis for overall survival as a function of age at diagnosis and residual disease after surgery in pre-chemotherapy patients, post-chemotherapy patients, and the entire cohort.

**Supplemental figure 5: Paired HGSOC tissue samples display senescence hallmarks following exposure to chemotherapy in patients.** (**a-f, n-o**) Matched pre- (left) and post-chemotherapy (right) tissue cores were stained in immunofluorescence for lamin B1 (**a**), Ki67 (**b**), IL6 (**c**), IL8 (**d**), vimentin (**e**), p16 (**f**), E-cadherin (**n**), and cleaved caspase-3 (**o**). Nuclei are counterstained with DAPI (blue). Epithelial cells are counterstained with cytokeratin 7, 18 and 19 (fuscia). (**g-m, p-q**) Quantification of the mean fluorescent intensity (MFI) of lamin B1 (**g**), Ki67 (**h**), IL6 (**i**), IL8 (**j**), vimentin (**k**), p16 (**m**), E-cadherin (**p**), and cleaved caspase-3 (**q**) in the epithelial (left) and stromal (right) tissue compartments of matched pre- and post- chemotherapy tissue cores. IL6, IL8, vimentin, E-cadherin, and cleaved caspase-3 MFI were quantified in the entire epithelial or stromal compartment, whereas lamin B1, Ki67, and p16 MFI were quantified in the nucleus. The red data points correspond to the cores used in the representative images in (**af, n-o**).

**Supplemental figure 6: Senescence-associated biomarker levels in post-chemotherapy HGSOC tissues correlate with 5-year survival.** (**a**) Table summarizing the direction of variation of different senescence markers after chemotherapy in the tumor epithelium and stroma, as well as their Kaplan-Meier log rank p-value comparing high and low post-chemotherapy expressors in terms of 5-year survival. Groups were separated into high and low expressors based on cut-offs determined by ROC analyses. H>L: high expressors have greater 5-year survival than low expressors; L>H: low expressors have greater 5-year survival than high expressors.

**Supplemental figure 7: MFI correlations between duplicate cores.** (**a**) Pearson correlations of the MFIs of different markers between duplicate cores in the epithelial (**a**) and stromal (**b**) compartments in the full set of cores on the TMA.

**Supplemental table 1: Summary of the characteristics of HGSOC primary cultures used for gene expression profiling:** Origins, passages in culture and E2F-response cluster classification.

